# Combination spinal cord stimulation at different frequencies produces sustained pain relief with immune activation

**DOI:** 10.64898/2026.08.06.742589

**Authors:** Yul Huh, Shaoyong Song, Toby Chen, Tianhe Zhang, Brad Hershey, Rosana Esteller, Ru-Rong Ji

**Affiliations:** Center for Translational Pain Medicine, Department of Anesthesiology, Duke University Medical Center, Durham, North Carolina, 27710, USA; Department of Cell Biology, Duke University Medical Center, Durham, North Carolina, 27710; Boston Scientific Neuromodulation Research and Advanced Concepts, 25155 Rye Canyon Loop, Valencia, CA 91355, USA; Department of Biomedical Engineering, University of Minnesota, Twin Cities, Minnesota, 55455; Department of Neurobiology, Duke University Medical Center, Durham, North Carolina, 27710; Department of Integrative Immunology, Duke University Medical Center, Durham, North Carolina, 27710

**Keywords:** Dorsal root ganglion (DRG), Innate immunity, neuroinflammation, rats, mice, RNA sequencing (RNAseq), spinal cord, spared nerve injury (SNI), spinal cord stimulation (SCS)

## Abstract

Spinal cord stimulation (SCS) is an established therapy for neuropathic pain, typically delivered at either low (60 Hz) or high (1 kHz) frequencies, with analgesic effects largely dependent on active stimulation. Here, we investigated whether combined-frequency SCS produces sustained analgesia beyond stimulation periods and explored the underlying mechanisms. Using a spared nerve injury (SNI) model in both rats and mice, we applied dual-frequency SCS (60 Hz + 1 kHz). This paradigm produced robust reversal of mechanical allodynia during stimulation and, notably, a progressive and long-lasting analgesic effect that persisted for days to weeks after stimulation cessation. RNA sequencing revealed pronounced immune-related transcriptional changes in the spinal cord, including upregulation of innate immune, pro-resolution, and neutrophil-associated pathways. Functional studies demonstrated that neutrophil depletion attenuated SCS-induced analgesia, whereas intrathecal S100A8 treatment mimicked therapeutic effects via CD69/SOCS3 signaling. These findings identify dual-frequency SCS as a promising strategy to prolong analgesia and highlight a critical role for neuroimmune modulation in sustained pain relief.

**Highlights:**

- Combined-frequency, not single-frequency SCS, sustains analgesia during washout
- Combination SCS induces robust immune activation in spinal cord and DRG
- Combination SCS increases spinal perfusion and promotes neutrophil recruitment
- Neutrophil signaling contributes to sustained SCS analgesia

## INTRODUCTION

More than 50 million Americans suffer from chronic pain^1^. The International Classification of Diseases (ICD) is widely used for the collection of public health data. The most common diagnostic categories in ICD-10 include: (1) other chronic pain, (2) low back pain, and (3) pain in limb; in ICD-11, the labeling categories are: (1) chronic cancer pain, (2) chronic peripheral neuropathic pain, and (3) chronic secondary musculoskeletal pain^2^. Low back pain (LBP), one of the most prevalent chronic pain conditions, encompasses a spectrum of pain types, including nociceptive, neuropathic, nociplastic, and non-specific pain. Consequently, LBP frequently overlaps with other pain conditions, such as neuropathic pain (e.g., sciatica)^3^. Neuropathic pain is a debilitating syndrome associated with pathological alterations in the structure, biochemistry, and function of both the peripheral and central nervous systems^3^. Increasing evidence suggests that neuroinflammation, characterized by infiltration of immune cells, glial activation, and the production of inflammatory cytokines and chemokines in the spinal cord and dorsal root ganglia (DRG), plays a crucial role in the development of neuropathic pain^4–7^. Furthermore, immune cells such as neutrophils, macrophages, and T cells can also facilitate neuropathic pain recovery and resolution through neuroimmune interactions ^8–12^.

Spinal cord stimulation (SCS) has been used to treat refractory chronic pain, such as LBP and neuropathic pain for several decades, resulting in pain relief in ~50% patients ^13^. Traditionally, lower frequency pulse trains (30-60 Hz) of SCS were used for pain management through activation of sensory afferents or their collaterals in the dorsal column ^13–16^, leading to “Gate Control” of the spinal cord pain circuit ^17–20^. Supraspinal mechanisms of SCS have also been proposed for the modulation of pain^16^. Recently, kilohertz (kHz) frequency pulse trains sub-perception SCS has shown efficacy in patients refractory to conventional medical management^21^. The lack of paresthesia and slower analgesia onset time ^22^ associated with kHz SCS suggest that additional mechanisms beyond the Gate Control Theory may also be attributable to the effects of SCS ^23,24^. It appears that kHz frequency SCS can be equally efficacious over a wide range of kHz frequencies, with 1 kHz SCS being the most energy-efficient in the 1-10 kHz range in a small double-blinded RCT ^25^. Furthermore, kHz frequency may not be always required, and a novel fast-acting sub-perception SCS therapy at 90 Hz was shown to produce a nearly 80% reduction in pain intensity at 3 and 6-months ^26^. Computational modeling has been used to assess the response of primary afferents, interneurons, and projection neurons to conventional, burst, and 10-kHz SCS. It was found that local cell thresholds were always higher than afferent thresholds and the recruitment order was the same ^27^. Thus, pain relief can be achieved by both paresthesia SCS at low frequencies and paresthesia-free SCS at kHz frequencies ^13^. In vivo recordings in rats revealed that spinal cord neurons including inhibitory neurons exhibited distinct responses to low and high frequency SCS ^28^. Thus, a combination of low and high frequency SCS may achieve additive and even synergistic benefits.

Mechanisms underlying SCS-induced pain relief remain incompletely understood. Notably, neuron-centered mechanisms alone cannot fully explain the sustained analgesia observed during the washout period. Recent studies have begun to investigate the effects of SCS on glial cells ^29–31^, and emerging evidence suggests that SCS may also modulate glial/immune cell activity in the spinal cord to regulate pain^32^. Understanding whether and how commercially available SCS therapies engage immune and glial mechanisms, and how to optimize these effects, may improve treatment options for chronic pain.

In the present study, using a rat spared nerve injury (SNI) model of neuropathic pain, we demonstrate that combined-frequency SCS (60 Hz + 1 kHz), but not single-frequency stimulation, produces sustained analgesia during the washout period. Furthermore, combined-frequency SCS induces robust immune activation in the spinal cord, suggesting a potential neuroimmune mechanism underlying its prolonged analgesic effects.

## RESULTS

### Combined-frequency SCS produces sustained relief of neuropathic pain in rats with SNI surgery

The present study was designed to compare the analgesic duration of low-frequency (60 Hz), high frequency (1 Kz), and combined-frequency (60 Hz + 1 kHz) SCS in rats using SNI model of neuropathic pain^33^ (Figure 1A and Supplementary Figure 1). Schematic of low frequency, high frequency, and dual frequency SCS. In our previous study, we examined the effects of single frequency SCS (1 x 6h) using a Boston Scientific device and observed only transient pain relief during the stimulation (Figure 1B) ^34^. In the current study, we modified the stimulation paradigm to four sessions of 2h each (4 x 2h), delivered every 48h, starting two weeks after SNI surgery. X-ray imaging confirmed the correct placement of the implanted electrodes (Figure 1B). Single-frequency SCS at either 1 kHz or 60 Hz elicited significant reduction of mechanical pain (allodynia), as indicated by increasing paw withdrawal threshold (PWT) in male rats; however, this effect was very transient, disappearing within 5h after stimulation (Figure 1C, D). Single frequency SCS also transiently alleviated cold hypersensitivity (duration of cold pain), with 1 kHz stimulation producing an earlier onset of analgesia compared to 60 Hz stimulation (Figure 1C, D).

**Figure 1.**
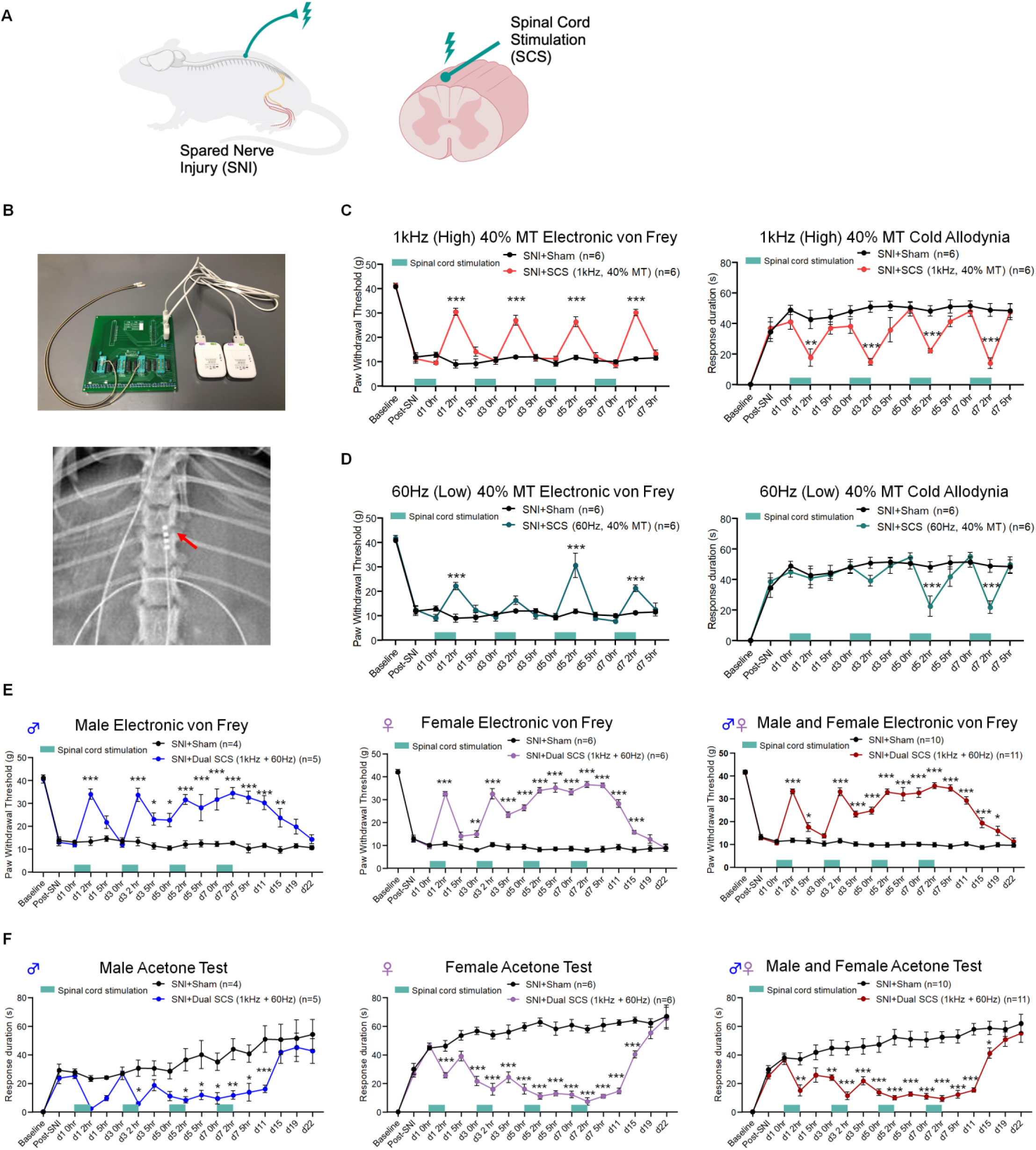
Combined-frequency SCS produces sustained relief of neuropathic pain in rats with SNI surgery. (A) Schematic of SNI surgery, electrode implantation, and SCS. (B) Top, SCS stimulator connected to rats with electrodes. Bottom, radiograph showing electrode implantation within the rat spinal column. The red arrow indicates the electrode contacts. (C, D) Effects of 1 kHz SCS (C) and 60 Hz SCS (D), given 2 h per day every other day for 4 sessions (indicated in green bards) at 40% of motor threshold. Left, Mechanical allodynia measured by paw withdrawal threshold (PWT) using electronic von Frey testing. Right, duration of cold pain (cold allodynia) in acetone testing. (E, F) Effects of combination frequency SCS (1 kHz and 60 Hz SCS) on mechanical pain (E) and cold pain (F) in male rats (left column, n=4 for control group, n=5 for SCS group), female rats (middle column, n=6), and males and females combined (right column, n=10 for control, n=11 for SCS). Electrode implantation was conducted 1 week after SNI and SCS was conducted approximately 2 weeks after SNI. All data expressed as mean ± SEM. \**P*<0.05, \*\**P*<0.01, \*\*\**P*<0.001, compared with control. Two-way ANOVA, followed by Bonferroni posthoc test.

Next, we tested whether analgesic effects would be enhanced or prolonged following combined-frequency SCS (simultaneous 60 Hz and 1 kHz). Strikingly, combined-frequency SCS produced a sustained analgesic washout effect following four stimulation sessions delivered over a 7-day period. This prolonged effect was observed for both mechanical allodynia (Figure 1E) and cold hypersensitivity (Figure 1F) across male, female, and combined-sex groups. During each 2-hour stimulation session, combined-frequency SCS fully reversed both mechanical and cold pain. Notably, a robust washout effect emerged during the inter-stimulation intervals and persisted after the fourth and final stimulation on day 7. The magnitude of this washout effect progressively increased with each successive stimulation session. Importantly, after the final stimulation, significant analgesia was maintained for an extended period, with mechanical pain relief lasting until day 19 and cold pain relief persisting until day 15 (Figure 1E, F).

### Combined-frequency SCS induces broad upregulations of innate immune and neutrophil-related genes

To investigate the mechanisms underlying sustained analgesia induced by combined-frequency SCS, we performed RNA sequencing of spinal cord tissues from SNI model rats following combined-frequency stimulation. Unexpectedly, RNA-seq revealed widespread upregulation of innate immune response genes, particularly those associated with neutrophil activity, in both the spinal cord and DRG of both sexes. RNA sequencing was conducted on whole spinal cord tissues collected from the lumbar enlargement of SNI rats implanted with electrodes and subjected to four 2-hour sessions of dual-frequency SCS. Heatmap analysis demonstrated broad gene upregulation compared to SNI rats receiving sham implantation (n = 3 male rats per group; Figure 2A). Gene Ontology (GO) analyses revealed significant enrichment of inflammatory processes, including leukocyte migration, regulation of inflammatory response, leukocyte cell–cell adhesion, myeloid leukocyte activation, and leukocyte chemotaxis (Figure 2B). Notably, many of the most highly upregulated genes were directly linked to neutrophil function, including *S100a8*, *S100a9*, *Mmp8*, *Mmp9*, *Ctsg*, *Mpo*, *Cxcl12*, *Ncf1*, *Defa5*, *Np4*, *Lcn2*, and *Serpinb1a* (Figure 2C).

**Figure 2.**
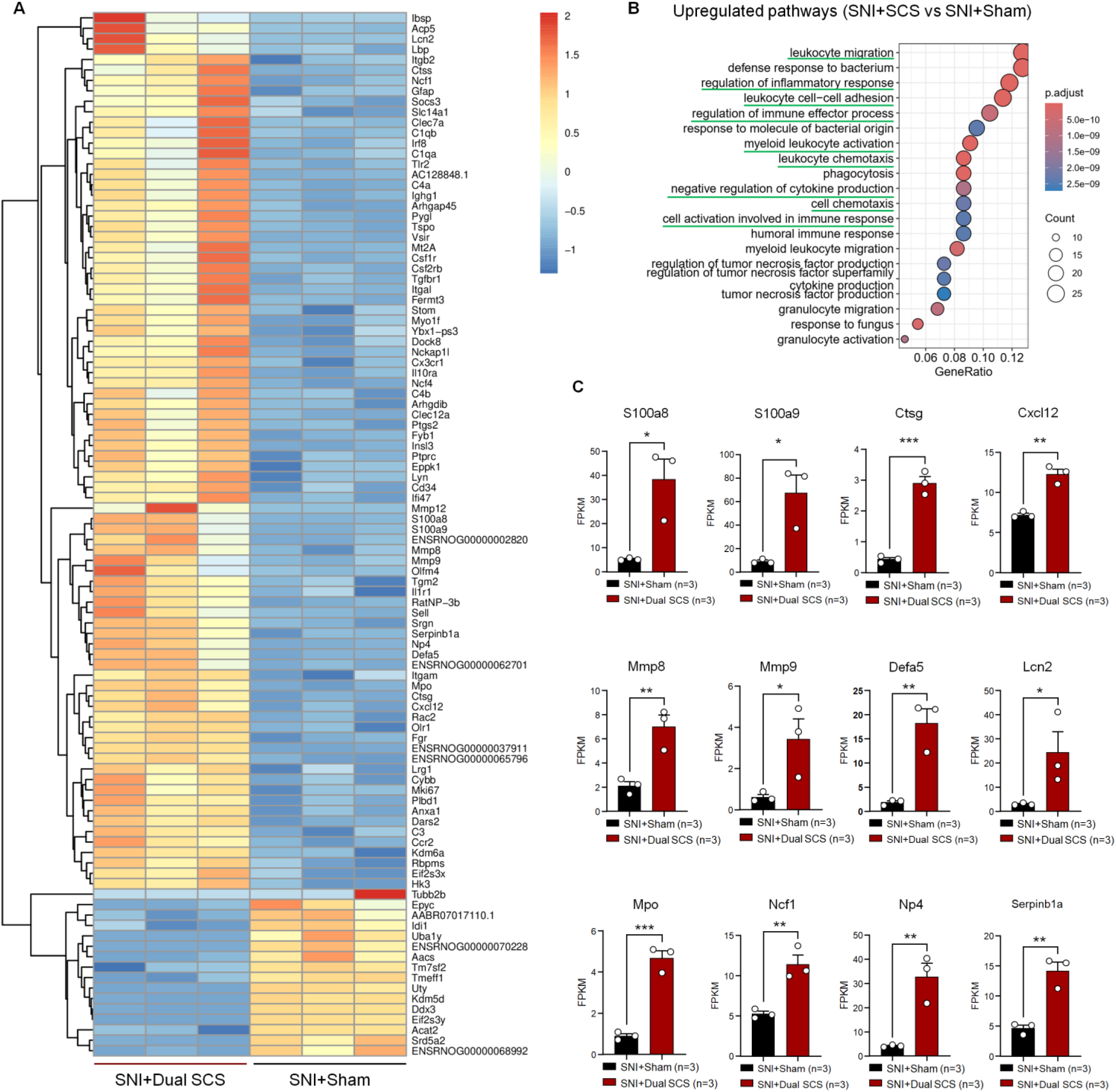
RNA-seq of spinal cord and DRG from dual-frequency SCS–treated SNI rats reveals broad upregulation of innate immune and neutrophil-related genes. (A) Heatmap of spinal cord gene expression comparing SCS-treated and SNI control rats. Tissues were collected after the fourth 2-hour SCS session (~2 weeks post-SNI). (B) GO analysis shows upregulation of immune processes, including leukocyte migration, inflammatory regulation, cell adhesion, myeloid activation, and chemotaxis. (C) Neutrophil-related genes are significantly upregulated in the spinal cord (FPKM values). n = 3 male rats per group. Data are mean ± SEM. Unpaired t-test (\**P*<0.05, \*\**P*<0.01, \*\*\**P*<0.001).

### Neutrophil signaling contributes to sustained analgesia following combined-frequency SCS

To determine whether neutrophils contribute to the analgesic effects of combined-frequency SCS, we administered SCS (40% motor threshold) to SNI rats following intrathecal injection of either an anti-polymorphonuclear neutrophil (anti-PMN) antibody or control rat IgG 1 hour prior to each of four 2-hour stimulation sessions. For mechanical sensitivity assessed by PWT, anti-PMN treatment did not affect the acute reversal of mechanical allodynia during the first and second SCS sessions (days 1 and 3). However, during the third session (day 5), the anti-PMN group exhibited a significantly reduced analgesic response compared to the IgG control group, with a further reduction observed during the fourth session (day 7). Importantly, depletion of neutrophils abolished the development of the SCS-induced analgesic washout effect. The anti-PMN group failed to exhibit sustained mechanical analgesia, displaying pain levels comparable to SNI rats receiving sham SCS. In contrast, the IgG control group maintained a robust washout effect similar to that observed with dual-frequency SCS alone (Figure 3A).

**Figure 3.**
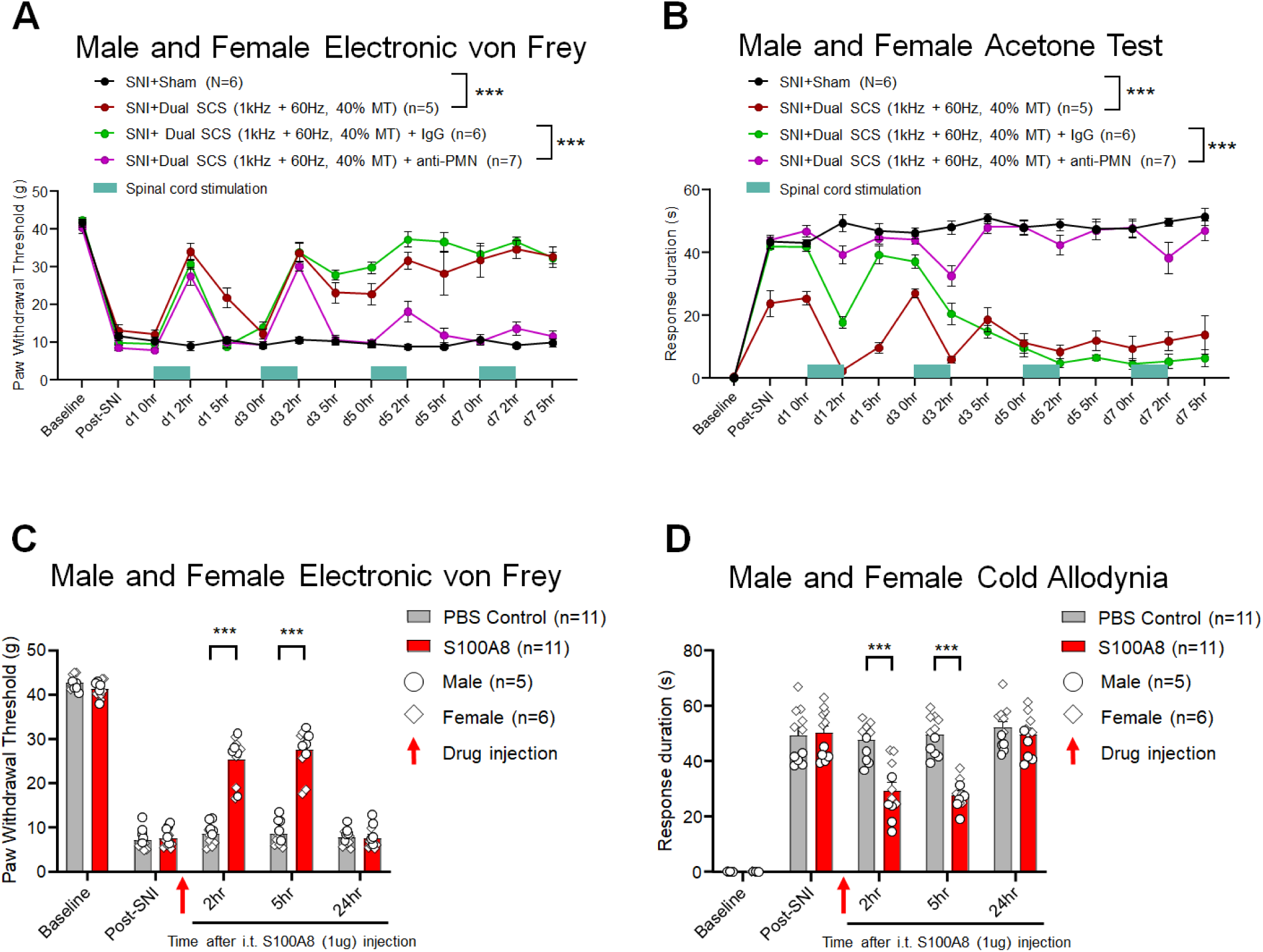
Neutrophil activity is required for sustained analgesia during the washout period after combined-frequency SCS. (A, B) Intrathecal anti-PMN (neutrophil-depleting) antibody blocks the analgesic washout effect of dual-frequency SCS. Anti-PMN (blue) or control IgG (green) was administered 1 h before each SCS session (days 1, 3, 5, and 7; ~2 weeks post-SNI). Mechanical (A) and cold (B) hypersensitivity were assessed. Anti-PMN treatment significantly attenuated sustained analgesia. (C, D) Intrathecal S100A8 (1 µg) reduces SNI-induced pain. S100A8 administration decreased mechanical (C) and cold (D) hypersensitivity at 2 and 5 h post-injection. Data are mean ± SEM. Statistical significance was determined by two-way ANOVA with Bonferroni post hoc testing. \*\*\**P*<0.001.

The results of mechanical pain testing were largely paralleled by cold allodynia assessed using acetone application to the hindpaw. Across all four SCS sessions, the anti-PMN group exhibited significantly greater cold allodynia compared to the control IgG group. Moreover, the SCS-induced cold analgesic washout effect failed to develop in the anti-PMN group, whereas the IgG control group displayed a robust and progressively enhanced washout effect with each successive SCS session (Figure 3B).

Given that S100A8 is a major cytosolic protein in neutrophils^35^, we next evaluated its analgesic effects by administering a single intrathecal injection of recombinant S100A8 to SNI rats. S100A8 treatment produced a significant reversal of mechanical allodynia at 2 and 5 hours post-injection (Figure 3C), as well as a marked reduction in cold allodynia at the same time points (Figure 3D) in male rats. These effects were transient, with pain behaviors returning to baseline levels by 24 hours, comparable to PBS-treated controls. Similar analgesic effects were observed in female rats (data not shown). These findings demonstrate that S100A8 exerts potent but transient analgesic effects in neuropathic pain.

RNA-seq analysis of spinal cord tissue collected 2 hours after intrathecal S100A8 (n = 4 males) versus PBS control (n = 4 males) revealed broad upregulation of immune-related genes, including *Socs3* and *Cd69* (Figure 4A, B). To further interrogate functional interactions, differentially expressed genes were analyzed using the STRING database and visualized via Sankey diagram. This analysis identified strong interactions among S100A8, SOCS3 (score: 0.26), and CD69 (score: 0.19), with convergence on MAPK1 signaling (score: 0.24) (Figure 4C).

**Figure 4.**
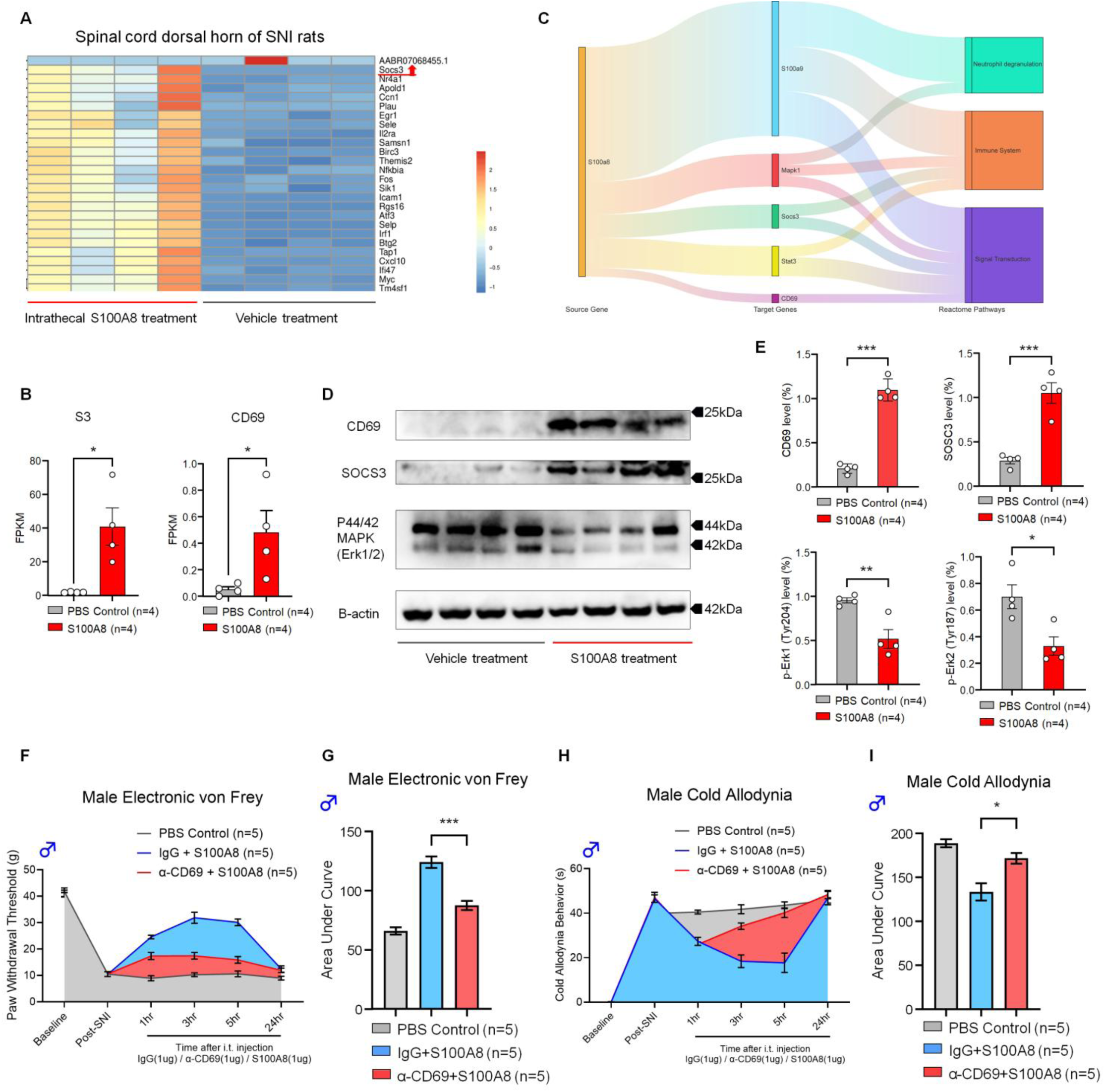
S100A8 reduces neuropathic pain via CD69/SOCS3/ERK signaling. (A) RNA-seq of spinal cord from SNI male rats treated with intrathecal S100A8 (2 h post-injection; n=4/group) versus PBS controls. Heatmap shows broad upregulation of immune-related genes. (B) RNA-seq quantification demonstrates significant increases in Socs3 and Cd69 expression (FPKM) following S100A8 treatment. (C) Interaction network of S100A8 with S100A9, MAPK1, SOCS3, STAT3, and CD69. (D–E) Intrathecal S100A8 increases CD69 and SOCS3 expression while reducing ERK1/2 phosphorylation. (D) Representative Western blots. (E) Quantification of CD69, SOCS3, pERK1, and pERK2 levels. (F–I) Blocking CD69 attenuates S100A8-induced analgesia in male animals. Intrathecal anti-CD69 antibody or control IgG was administered 1 h before S100A8. Mechanical (F, G) and cold (H, I) hypersensitivity were assessed at 1, 3, 5, and 24 h. AUC analysis shows reduced analgesic efficacy with CD69 blockade. Data are mean ± SEM. Statistical significance: unpaired t-test (B, E) and one-way ANOVA with Bonferroni post hoc test (G, I). \**P*<0.05, \*\**P*<0.01, \*\*\**P*<0.001, \*\*\*\**P*<0.0001.

Among the most significantly upregulated genes was *Socs3*, along with its upstream-associated receptor *Cd69* (Figure 4B). These changes were validated at the protein level by Western blot, showing increased SOCS3 and CD69 expression in the spinal cord following S100A8 treatment (Figure 4D, E). Nerve injury is known to enhance ERK phosphorylation in the spinal cord, contributing to neuropathic pain^36,37^. Notably, S100A8-induced analgesia was associated with marked inhibition of phosphorylated ERK1/2 (pERK1/2) (Figure 4D, E).

To determine whether CD69 mediates S100A8-induced analgesia, we pretreated SNI rats with intrathecal anti-CD69 antibody or control IgG prior to S100A8 administration. In von Frey testing, anti-CD69 pretreatment significantly reduced the analgesic effect of S100A8, as indicated by a decreased area under the curve (AUC) for PWT across 1, 3, 5, and 24 hours (Figure 4F, G). Similarly, cold allodynia testing revealed significantly increased AUC values in the anti-CD69 group, indicating reduced analgesia (Figure 4H, I).

Together, these results demonstrate that S100A8-induced analgesia is mediated, at least in part, through CD69-dependent signaling pathways.

## DISCUSSION

Despite the clinical benefits of SCS for chronic pain, there remains a limited understanding of its underlying mechanisms, particularly with the increasing use of high frequency (kHz range) SCS. Traditionally, SCS analgesia has been explained by the Gate Control Theory, whereby activation of large-diameter, myelinated Aβ fibers inhibits nociceptive transmission from small-diameter C fibers in the dorsal horn-fibers^23^. However, accumulating evidence suggests that high-frequency SCS engages additional mechanisms beyond gate control. Notably, clinical studies report the absence of paresthesia and a delayed onset of analgesia with kHz-frequency stimulation, indicating distinct underlying processes^22^.

In this study, we demonstrate that combined-frequency SCS (60 Hz + 1 kHz) generates a robust and extended analgesic washout effect in the rat SNI model of neuropathic pain. This prolonged analgesia cannot be readily explained by neuron-centered mechanisms alone and instead points to the involvement of non-neuronal pathways. Emerging evidence has implicated glial cells in SCS-induced analgesia^29,30^. Recently, neutrophil activation was found to promote the resolution of inflammatory pain^35^. Consistent with this framework, our findings reveal that combined-frequency SCS induces profound immune modulation in the spinal cord, which contributes to sustained analgesia during the washout period. RNA-seq analysis demonstrated broad upregulation of innate immune and neutrophil-related genes, and functional studies support a critical role for neutrophils in mediating this effect. Notably, depletion of polymorphonuclear neutrophils abolished the development of the analgesic washout effect.

Mechanistically, we identify neutrophil-derived S100A8 as a key mediator of this process. Intrathecal administration of S100A8 produced potent, albeit transient, analgesia and activated a SOCS3/CD69-dependent signaling pathway in the spinal cord. This pathway was associated with suppression of ERK phosphorylation, a known contributor to central sensitization^37^. Furthermore, blockade of CD69 significantly attenuated S100A8-induced analgesia, supporting a functional role for this signaling axis.

Although immune activation is classically associated with pain hypersensitivity, particularly through microglia, astrocytes, and macrophages ^6,38–41^, recent studies highlight a dual role of the immune system in both the initiation and resolution of pain ^35,42^. Furthermore, immunotherapy has been proposed for the treatment of chronic pain ^43,44^. In this context, our findings suggest that SCS can harness pro-resolving immune mechanisms to achieve sustained analgesia. These results provide a conceptual framework in which neuromodulation engages neuroimmune interactions, shifting the spinal cord microenvironment toward a pain-resolving state.

In summary, our study identifies a previously unrecognized neuroimmune mechanism underlying SCS-induced analgesia and suggests that targeting neutrophil signaling and S100A8-mediated pathways may enhance the therapeutic efficacy of SCS for chronic pain.

## RESOURCE AVAILABILITY

### Lead contact

Further information and requests for resources should be directed to and will be fulfilled by the lead contact, Ru-Rong Ji.

### Materials availability

This study did not generate new unique reagents.

## ACKNOWLEDGEMENTS

The work was supported by Duke University Anesthesiology Research Funds and a grant from Boston Scientific (R.-R.J). This study was also supported by the NIH grant R01NS13182 and DoD grants W81XWH2110756 and W81XWH2210646 to R.-R.J.

## AUTHOR CONTRIBUTIONS

Y.H. conducted spinal surgery, SCS, behavioral, RNAseq experiments in rats. S.S conducted SCS in mice and assisted with single-cell data analysis. R.-R. J. supervised the project. T.Z., B.H., and R.E. participated in the project discussion. Y.H and R.-R.J. wrote the paper, and all the co-authors edited and approved the manuscript.

## DECLARATION OF INTERESTS

T.Z. and B.H. are employees of Boston Scientific Inc., and R.E. was an employee of Boston Scientific when the work was performed. T.Z. holds Boston Scientific stock.

## SUPPLEMENTAL METHODS

### Animals

Male and female Sprague-Dawley rats (200-250 g) and CD1 mice (8-12 weeks) of both sexes were obtained from Charles River Laboratories. Animals were housed in randomly assigned sex-matched pairs on a 12-hour light and 12-hour dark cycle at a room temperature of 22 degrees Celsius with ad libitum food and water. All animal experiments in this study followed the National Institutes of Health Guide for the Care and Use of Laboratory animals. The Institutional Animal Care Use Committee (IACUC) of Duke University approved this study prior to the start of experiments.

### Spared nerve injury model of neuropathic pain

Spared Nerve Injury (SNI) surgery was done on the sciatic nerve in rats, with the tibial and common peroneal nerves ligated with a 5-0 suture and then transected and then a 3-5 mm section was removed as detailed previously ^33,34^. The sural nerve was carefully left undisturbed during the surgery. Resulting neuropathic injury was measured by von Frey filament and acetone application to the lateral aspect of the ipsilateral hindpaw corresponding to the dermatome region innervated by the sural nerve.

### Spinal cord stimulation (SCS) electrode implantation

SCS electrode implantation was conducted one week after post-SNI surgery. Laminectomy was performed at the T10-T12 vertebral level with a microdrill, and a cylindrical four-contact electrode was introduced into the epidural space in a rostral orientation. The electrode wiring was fixed outside the laminectomy opening by suture to the surrounding muscle, and the remaining wire was routed beneath the skin to the base of the neck to connect with the interface adapter. An external neuro-stimulator device (Boston Scientific Wavewriter External Trial Stimulator) was connected to the adapter by cable and programmed via wireless connection to a Windows tablet PC provided by Boston Scientific. Motor threshold (MT) was measured by gradual increase of current amplitude until muscle contractions were observable under 2% inhaled isoflurane.

### Spinal cord stimulation settings and treatment in rats

SCS treatment was started 7 to 14 days after electrode implantation (2 to 3 weeks after SNI surgery). Animals receiving single frequency SCS received either 1 kHz “high frequency” pulse trains or 60 Hz “low frequency” pulse trains at 40% motor threshold. Animals receiving dual frequency SCS received simultaneous 1 kHz and 60 Hz symmetric biphasic stimulation at 40% motor threshold. The stimulation treatment regimen consisted of four total 2-hour SCS treatments each given 48 hours apart (days 1, 3, 5, and 7 at 0h timepoint). Stimulation was conducted on animals while fully awake and freely moving in individual enclosed plastic boxes on elevated mesh metal flooring at stable room temperature and humidity.

### Behavior tests for mechanical and cold sensitivity

Rats were habituated in plastic boxes for 2 days prior to baseline behavior testing. Mechanical sensitivity was measured by assessment of paw withdrawal threshold (PWT) using an electronic von Frey anesthesiometer (IITC Life Science Inc.) with a filament force range of 0 to 50 g when applied to the lateral aspect of the hind paw. Cold sensitivity was measured by application of 150 microliters of acetone by micropipette to the lateral aspect of the hind paw. The duration in seconds of cold allodynia behavior, including lifting, licking, and guarding of the paw over a 90 second period was measured. All behavioral tests were conducted in a blinded manner.

### RNA isolation and sequencing

Total RNA was extracted from the rat lumbar spine cord and dorsal root ganglion using the Trizol reagent (Thermo Fisher Scientific, Cat No.15596018) according to the manufacturer’s protocol. Temporarily, these tissues were homogenized in Trizol reagent (0.1g/1ml) and incubated at room temperature for 5 minutes. Next, 0.2 mL of chloroform (Thomas Scientific, CAS No.67-66-3) was added, and the mixture was shaken for 15 seconds, followed by incubation at room temperature for 5 minutes. The sample was centrifuged at 12,000 rpm for 15 minutes at 4°C. The aqueous phase was collected, and RNA was precipitated with an equal volume of isopropyl alcohol (Sigma-Aldrich, CAS No.67-63-0), incubated at room temperature for 10 minutes, and then centrifuged at 12,000 rpm for 10 minutes at 4°C. The RNA pellet was washed with 70% ethanol and transferred to a 2ml collection tube and centrifuged for 15 s at ≥8000 × g. After removing the ethanol, 700 μl Buffer RW1 (QIAGEN, Cat.No.74104) was added to the RNeasy spin column and centrifuged for 15 s at ≥8000 × g and discarded the flow-through. Then, 500 μl Buffer RPE (QIAGEN, Cat.No.74104) was added to the RNeasy spin column and centrifuged for 15 s at ≥8000 × g and discarded the flow-through. Next, we added 500 μl Buffer RPE (QIAGEN, Cat.No.74104) to the RNeasy spin column, centrifuged for 2min at ≥8000 × g and discarded the flow-through. After placing the RNeasy spin column in a fresh 1.5 ml collection tube, RNase-free water was added directly to the center of the membrane. The column was left alone for 1 minute, then centrifuged at the recommended speed to elute the RNA. RNA concentration and purity were assessed using a NanoDrop spectrophotometer (Thermo Fisher Scientific) and an A260/280 ratio between 1.8 and 2.0. RNA integrity was confirmed by Agilent Technologies 2100 Bioanalyzer (Agilent Technologies) to assess RNA Integrity Number (RIN) values > 7.0 as a threshold for high-quality samples used in Poly (A) RNA sequencing library preparation. The Poly (A) RNA Sequencing library was generated by using the TruSeq RNA preparation kit (Illumina, Cat.No. RS-122-2001, RS-122-2002), following Illumina’s TruSeq-stranded-mRNA sample preparation protocol. In brief, the mRNA was purified by Oligo (dT) magnetic beads with two rounds of purification. Then, poly (A) RNA was fragmented into 200 bp short fragments using divalent cation buffer in elevated temperature. Fragmented RNA was then reverse transcribed into cDNA using random hexamer primers and cDNA synthesis kit. Double-stranded cDNA was synthesized, end-repaired, and A-tailed to prepare for adapter ligation. The concentration of cDNA was assessed by using the Qubit 2.0 (Thermo Fisher Scientific), and the length of library fragments was determined by using the Agilent 2100 Bioanalyzer. Finally, Paired-ended sequencing was performed by using an Illumina’s NovaSeq 6000 platform (Illumina).

### Differential expression analysis of mRNAs

Fragments Per Kilobase of transcript per Million mapped reads (FPKM) was used for the normalization and calculation of mRNA expression level by StringTie, and the R package DESeq2 (http://bioconductor.org/ packages/DESeq2/) was utilized for filtering the differentially expressed genes (DEGs). The raw *p*-value was adjusted to false discovery rate (FDR), and FDR below 0.05 and absolute fold shift ≥ 2 were considered differentially expressed mRNAs. To summarize the characteristics of gene expression profiles, volcano plots and heatmaps were created using the R package *ggplot2*.

### GO enrichment analysis of differentially expressed genes

Gene ontology (GO) (http://www.geneontology.org) is a major bioinformatics initiative that provides structured, controlled vocabulary for describing the molecular functions, cellular components, and biological processes associated with gene and gene product across all species. We utilized the R package *clusterProfiler* to get GO enrichment analysis of DEGs. The adjusted *p*-values (p < 0.05) was set as the threshold value. The results were visualized using *barplot()*, *dotplot()*, and *emapplot()* for intuitive presentation.

### KEGG enrichment analysis of differentially expressed genes

KEGG (Kyoto Encyclopedia of Genes and Genomes) (http://www.kegg.jp/) is a collection of databases dealing with genomes, biological pathways, diseases, drugs, and chemical substances. We used *enrichKEGG()* in the R package *clusterProfiler* to perform KEGG pathway enrichment analysis for DEGs. The adjusted *p*-value cutoff was set at 0.05 for statistical significance. *barplot*, *dotplot*, and *emapplot* are visualization functions for enriched KEGG pathways.

### Western blot analysis

We used western blot analysis to validate the expression changes in SOCS3, CD69, and the MAPK1 pathway and assess protein level alterations associated with the interactions and pathway involvement identified in the RNA sequencing and STRING database analysis. Spinal cord tissues were homogenized in RIPA lysis buffer (Sigma, Cat No. R0278) containing a protease inhibitor cocktail tablet (pH 7.4) (Roche Diagnostics, Cat No. 5892988001) and phosphatase inhibitor cocktail (Cell Signaling Technology, Cat No.5870S). The protein concentration was measured via bicinchoninic acid (BCA) protein assay kit (Thermo Fisher Scientific, Cat No.23227). The supernatant was mixed with 5x SDS-PAGE protein loading buffer (Boster, Cat No. BST19F28C12) containing β-mercaptoethanol (BME) for reduction, then boiled for 5 minutes at 95°C to prepare for Bis-Tris PAGE. Protein samples were separated on 10% SDS–PAGE gels at 120 V for 1 h and transferred onto a 0.2 μm polyvinylidene fluoride (PVDF) membrane (BioRad, Cat No.1620177) at 200 mA, 4°C for 1.5 h. The membrane was blocked with blocking buffer at room temperature for 2 hours and incubated with the following primary antibodies at 4°C overnight: CD69 polyclonal antibody (Invitrogen, Rabbit, 1:1000, Cat No. PA5-114989), SOCS3 polyclonal antibody (Invitrogen, Rabbit, 1:500, Cat No. PA5-87485), Phospho-p44/42 MAPK (Erk1) (Tyr204)/(Erk2) (Tyr187) antibody (Cell Signaling Technology, mouse, 1:2000, Cat No. 5726S), and beta actin polyclonal antibody (Proteintech, Rabbit, 1:2000, Cat No. 20536-1-AP). The membrane was washed with TBST (5 washes, 5 minutes each) and incubated with horseradish peroxidase-conjugated secondary antibodies: anti-rabbit IgG, HRP-linked antibody (Cell Signaling Technology, 1:2000, Cat No. 7074P2), and mouse anti-rabbit IgG-HRP antibody (Santa Cruz, 1:2000, Cat No. sc-2357) for 2 h at room temperature. Protein bands were visualized by Azure Imaging Systems (Azure Biosystems) with the HRP substrate (Millipore, Cat No. WBLUR0100). The protein expression was normalized to β-actin control.

### Statistics

The data presented in this study were expressed as the mean ± the standard error of measurement. Statistical analyses were conducted with Prism 10.2.3 software (Graphpad). Behavioral data were amazed by two-tailed student’s t-test, one-way or two-way ANOVA and Bonferroni’s post-hoc test. Statistical significance was measured at *p* < 0.05.

