## Supplemental Fig 1 for "Combination spinal cord stimulation at different frequencies produces sustained pain relief with immune activation"

### Supplementary Figures: 1

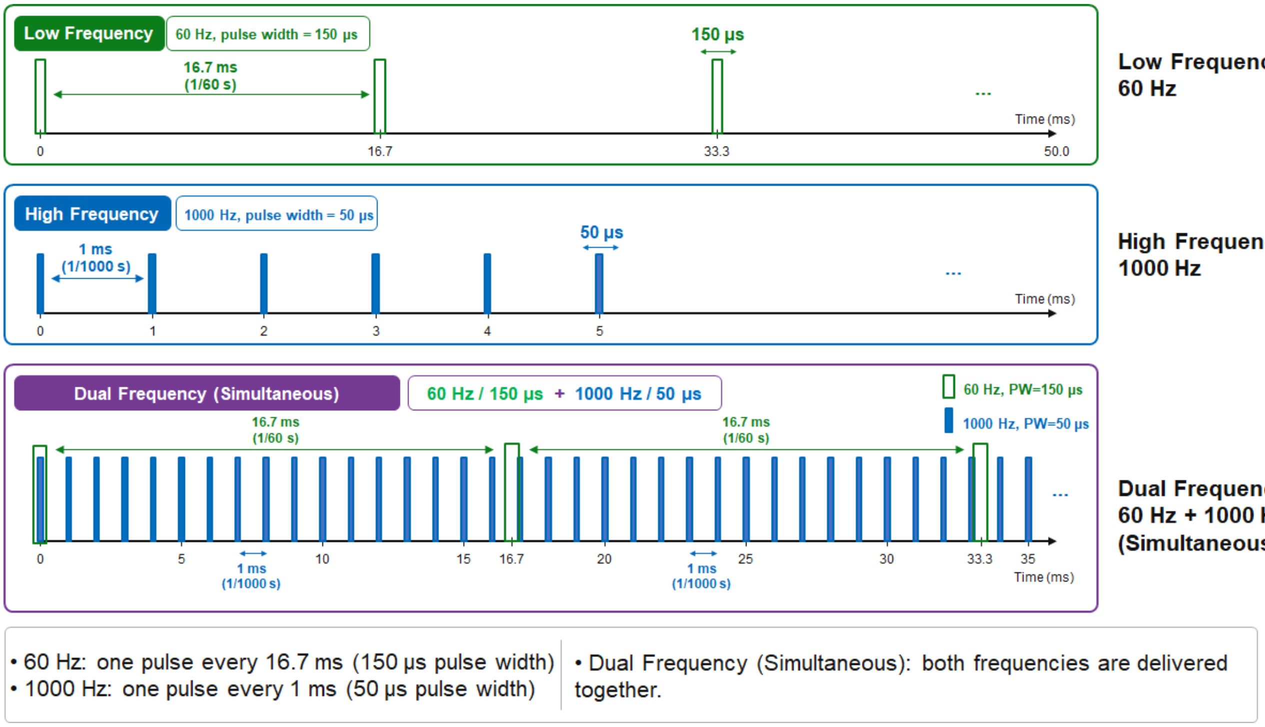


**Supplementary Figure 1**. Schematic of low frequency, high frequency, and dual frequency SCS.
